# Migration and establishment of progenitor pool of melanocytes is governed by SEMA3E-PLXND1 signaling

**DOI:** 10.1101/2023.02.28.530560

**Authors:** Yogaspoorthi Subramaniam, Babita Sharma, Ayush Aggarwal, Desingu Ayyappa Raja, Iti Gupta, Madeeha Ghazi, Sridhar Sivasubbu, Vivek T Natarajan

## Abstract

Vertebrate pigmentation is an outcome of an interplay of several signaling pathways that result in immense diversity of pigment patterns observed across the animal kingdom. Transitory nature of these signaling events impedes deciphering pathways that control migration and establishment of melanocyte stem cells (McSC), necessary for pigment patterning. Using zebrafish and cultured mammalian melanocytes, we uncover a hitherto unknown role for Plexin D1 signaling. This pathway directs migration by F-actin modulation and further dictates subsequent functional states of melanocytes through a transcriptional response. In zebrafish, abrogation of PLXND1 derails melanocyte migration and reduces mid-line melanophores emerging from the regeneration competent McSC pool. In cultured melanocytes, activation of PLXND1 by the ligand semaphorin 3E reduces the velocity of migration, influences directional correlation, and promotes movement towards positive cues such as SCF. PLXND1 activation results in EGFR signaling necessary for McSC establishment, and induces GNAS, an effector of MC1R pathway involved in melanocyte maturation. Identification of this long-range secreted negative chemotactic signaling provides a missing player and enriches reaction diffusion model for pigment patterning.

## Introduction

Migration of neural crest cells (NCC) is accompanied with concomitant specification and differentiation leading to generation of melanocytes among other cell types [1]. It is likely that the spatial organization of the chemotactic microenvironment influences the migratory path and timely acquisition of the final differentiated state. Thereby, understanding the permissive and restrictive embryonic landscapes that regulate NCC streaming gains importance [2]. Several signaling pathways including SCF, WNT3A, BMP4, ET1 *etc* are known to control the formation of melanocytes, and also influence their migration [3-9]. Among these the central role played by the cKit-Kitlg signaling (SCF or stem cell factor in mammals), in melanocyte migration is well established [4, 10].

The organized streaming of melanocyte precursors establishes Melanocyte stem cell (McSC) pool in defined areas such as hair follicle bulge region in mammals and near the dorsal root ganglion in zebrafish. The establishment of McSC is dependent on ERBB (an EGFR family receptor) and KIT signaling. Both genetic as well as pharmacological inhibition of ERBB signaling abrogates McSC establishment [11]. These McSCs are important for late-stage zebrafish larval melanocytes present in the lateral mid-line, as well as regeneration of pigmentation during injury and metamorphosis [11]. However, the mechanistic details of the stem cell pool establishment as well as its recall during regeneration remains elusive.

During migration, the transiting path of NCCs is restricted to well-defined routes by inhibitory signals [12]. Most prominent among the repulsive interactions is the contact inhibition of locomotion between migrating neural crest cells [13]. Additionally, several classes of Semaphorins and their cognate receptor Plexins are known to guide migratory cells by providing environmental cues. Detailed understanding of cellular pathfinding comes from the study of neuronal synaptic assemblies. Herein vertebrate-specific Semaphorins of the class 3 restrict migration of neuronal growth cones by activating both Plexins and Neuropilins (Nrps/Npns) [14, 15]. Similarly, SEMA6-PLXND1 axis is involved in navigating the cardiac NCCs during early vertebrate cardiac development [16]. SEMA3E-PLXND1 axis coordinates axonal extension and steering, in the developing nervous system [17]. Previously, this signaling is known to play a crucial role in branching and pathfinding morphogenesis of blood vessels during cardiovascular development (reviewed in [18]). Secreted inhibitory cues have been known, but so far, they have not been elucidated for melanocyte migration.

In this study, we analysed the responsiveness of melanocyte precursors as they migrate across defined embryonic landscapes by studying the transcriptional regulation of cell surface receptors for guidance cues at different stages of melanocyte development. Plexin-D1 - semaphorin-3E signaling emerged as a pathway that controls melanocyte migration by the focal alterations of actin-cytoskeleton. Further this pathway cross-talks with EGFR signaling and modulates the establishment of melanocyte stem cell pool in zebrafish. Transcriptional activation brought about by this pathway preconditions cells to melanocortin signaling by upregulating GNAS, that promotes melanocyte differentiation. By identifying a key regulator of melanocyte behaviour our study provides crucial insights into the pigment pattern formation.

## Results

### *plxnd1* is essential for melanophore patterning during development in zebrafish

Receptor expression on cell surface is indicative of the cell’s functional state. As NCCs progressively differentiate into patterned melanocytes, the expression levels of receptors that regulate biochemical and morphological properties are likely to change. To understand the basis of melanocytes developmental fate we isolated GFP expressing melanocytes from the transgenic *Tg (mitfa:GFP*) line during consecutive stages of their differentiation and analysed the expression of surface receptors and associated biological processes (Fig 1A and Supplementary Fig S1A-C).

**Fig 1:**
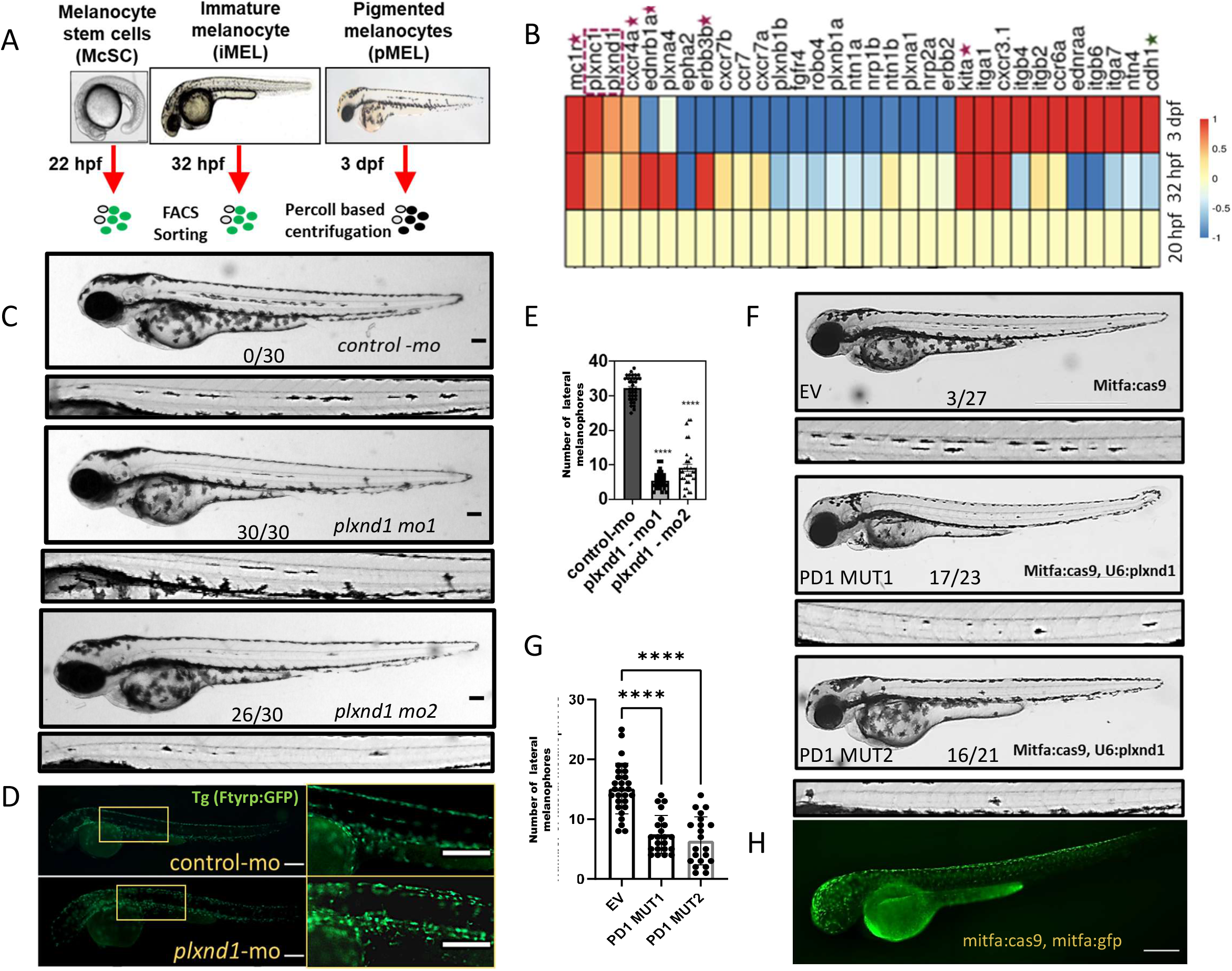
plxnd1 is required for lateral melanophore pattern establishment during early development. (A) Schematic representing experimental strategy for expression analysis of cell surface receptors during progressive stages of melanocyte development. (B) Expression heat map of differentially regulated cell surface receptors. Receptors with known functional roles in melanocyte development (magenta star). (C) Lateral embryonic melanophore patterning in plexin receptor morphants at 3dpf. Numbers represent embryos with <10 lateral melanophores out of total scored in one biological replicate out of three. Scale bar = 100µm. (D) Lateral embryonic melanophore patterning in plxnd1 MO 1 and control MO in Tg(fTyrp1: GFP) fishes that mark differentiating melanophores at 48 hpf. (E) Quantitation of lateral line melanophore in morphants (Fig 1C) at 3dpf. error bar: mean ±s.e.m across 3 biological replicates. p-value<0.001 is depicted by ***. One way ANOVA Dunnett’s multiple comparisons test. (F) f0 zebrafish embryos at 3 dpf injected either with plasmid driving mitfa:Cas9, mitfa:GFP or with mitfa:Cas9, mitfa:GFP, U6:plxnd1 sgRNA1 and 2. Zoomed insets of lateral line melanophores is depicted at the bottom. (G) Quantitation of lateral line melanophores in mutants represented in (Fig 1F) at 3dpf. error bar: mean ±s.d. p-value<0.0001 is depicted by ****. Scale bar = 100µm. One way ANOVA Dunnett’s multiple comparisons test. (H) 2 dpf zebrafish embryos injected with mitfa:Cas9, mitfa:GFP plasmid.

Like most NCCs, melanocyte precursors adhere to strict routing pattern within the designated dorso-ventral and medial streams [11]. In zebrafish both these streams originate almost simultaneously [19]. The component of directional persistence during such migration is critical. Consistent with this, most chemotactic receptors including robos, netrins, ephrins and plexins were found differentially regulated during this developmental window (Fig 1B and Supplementary Fig S1B). While, majority of guidance receptors were drastically downregulated during melanocyte pattern development, members of plexin family namely, *plxnc1* and *plxnd1* were found to be progressively increased in expression (Fig 1B). Based on studies from hair follicle profiling, plexin C1 is known to have a prominent role in melanocyte stem cell maintenance [20]. Other class of plexin receptors, *plxnb1a, plxnb1b* and *plxna4* were found to be significantly decreased in expression in pigmented melanophores. The role of these plexin receptors in melanocyte migration if any, isn’t known so far.

To delineate the role of plexin receptors in melanocytes we performed a quick screen using anti-sense morpholino based gene silencing strategy and analysed pigment patterning in developing zebrafish embryos. Morpholino-based silencing of *plxnd1* led to significant loss of pigmented melanophores at lateral mid-line with respect to the control embryos at 72 hours post fertilisation (hpf) (Fig1C & ID). Whereas, MO targeting other Plexins (*plxnb1a, plxnb1b, plxna4*) did not show this phenotype at the highest permissible concentration where viability was close to 85%. Even at very low doses, *plxnc1* morphants were lethal, hence its role if any in lateral mid-line formation could not be assessed (Supplementary Fig S1D&E). The *plxnd1* morpholino used herein (*plxnd1* MO1) targets splicing and has been reported earlier [21]. Another sequence independent morpholino targeting the ATG site (*plxnd1* MO2) resulted in a similar reduction in lateral mid-line melanophores (Supplementary Fig S1D). Using a custom synthesised antibody, we could demonstrate a reduction in the level of *plxnd1* (Supplementary Fig S1F). CRISPR-based global ablation of *plxnd1* in zebrafish using ribonucleoprotein complex also resulted in a similar phenotype (Supplementary Fig S1H&I). While the lack of involvement of *plxnb1a, plxnb1b* and *plxna4* on lateral mid-line melanophore establishment is difficult to comment solely on the basis of this morpholino screen, the phenotype of reduction in lateral mid-line melanophores appears to be selective to silencing of *plxnd1*.

Previous studies demonstrate genetic ablation of *plxnd1* in zebrafish leads to defects in vasculature and corresponding studies in mice suggest *plxnd1* ablation to be prenatally lethal [22]. While the global CRISPR mutants were alive during the first few days, there was a marked lethality. Thereby, we adapted a cell-type specific knockout strategy using MinicoopR vector (addgene118844) created in Leonard Zon’s lab. This plasmid drives Cas9 and *mitfa* mini gene under *mitfa* promotor (specific to melanocytes) and has a space for two sgRNA under ubiquitous promotor [23]. We replaced *mitfa* mini gene by GFP using restriction digestion approach to mark the melanophores (Fig 1H). Using this plasmid melanophore specific ablation was attempted using two different sequence independent sgRNA targeting *plxnd1*, while the control involved a non-targeting sgRNA. This strategy also addresses the cell autonomic functions of *plxnd1* and prevents phenotypes of *plxnd1* in other cell types from confounding our interpretations. With this strategy we recapitulated the lateral melanophore reduction phenotype (Fig 1F). Thereby using three independent approaches, namely two sequence independent morpholino based silencing, two sequence independent global knockout and two sequence independent melanocyte specific knockout, we demonstrate the selectivity of *plxnd1* in establishing the lateral mid-line melanophores.

Using transgenic melanocyte reporter line Tg(f*Tyrp1*: GFP), we observed that early patterning of embryonic melanocytes at 48 hpf was substantially altered in *plxnd1* morphants (Fig 1D). The aberrant positioning of immature melanocytes at 48hpf could have led to defective embryonic pattern development at 3 days post fertilisation (dpf). Thereby we conclude that *plxnd1*is required for populating the lateral line melanophores. In our earlier study involving pigmenting melanocyte associated gene expression changes, we had observed *Plxnd1* to be regulated during the pigmentation program of cultured mouse B16 melanoma cells [24, 25], suggesting an important role of *plxnd1* in melanocyte during pigmentation. Though other plexins could have melanocyte specific functions, in this study we focus on investigating this signaling pathway further in detail.

### Loss of *plxnd1* leads to delay in melanocyte precursor streaming and aberrant embryonic stripe pattern formation

The *plxnd1* morphants show apparently normal distribution of melanocytes and only the lateral mid-line melanocytes are severely reduced. A likely possibility could be the improper migration of immature melanocyte precursors to home in the dorsal root ganglion and establish the McSC pool. We therefore performed time-lapse imaging of *Tg (mitfa:GFP)* labelled zebrafish morphants to track melanocyte migration. Streaming of these GFP-tagged cells proceeds in an orchestrated manner. Migrating melanocytes in zebrafish appear along both the neural crest streams: dorso-lateral and ventral, almost simultaneously and then segregate into lateral stripes (Fig 2A). In control morphants between 20-32 hpf active melanocyte migration was clearly observed (Supplementary Video V1). Melanocytes were found both populating the dorsal stripe, migrating towards the posterior end, and arising into ventral streams. However, in *plxnd1* morphants the GFP-tagged cells were found to be restricted at the dorsal end. Both streams of migration were significantly inhibited (Supplementary Video V2). While melanocyte streams were clearly visible at dorsal trunk region at 20 hpf, their appearance towards the posterior regions leading to tail was defective in *plxnd1* morphants (Fig 2B). More strikingly *plxnd1* morphants had far fewer streams of ventrally migrating melanocytes (Fig 2C) that migrate to a shorter distance compared to the streams originating in control morphants (Fig 2D).

**Fig 2:**
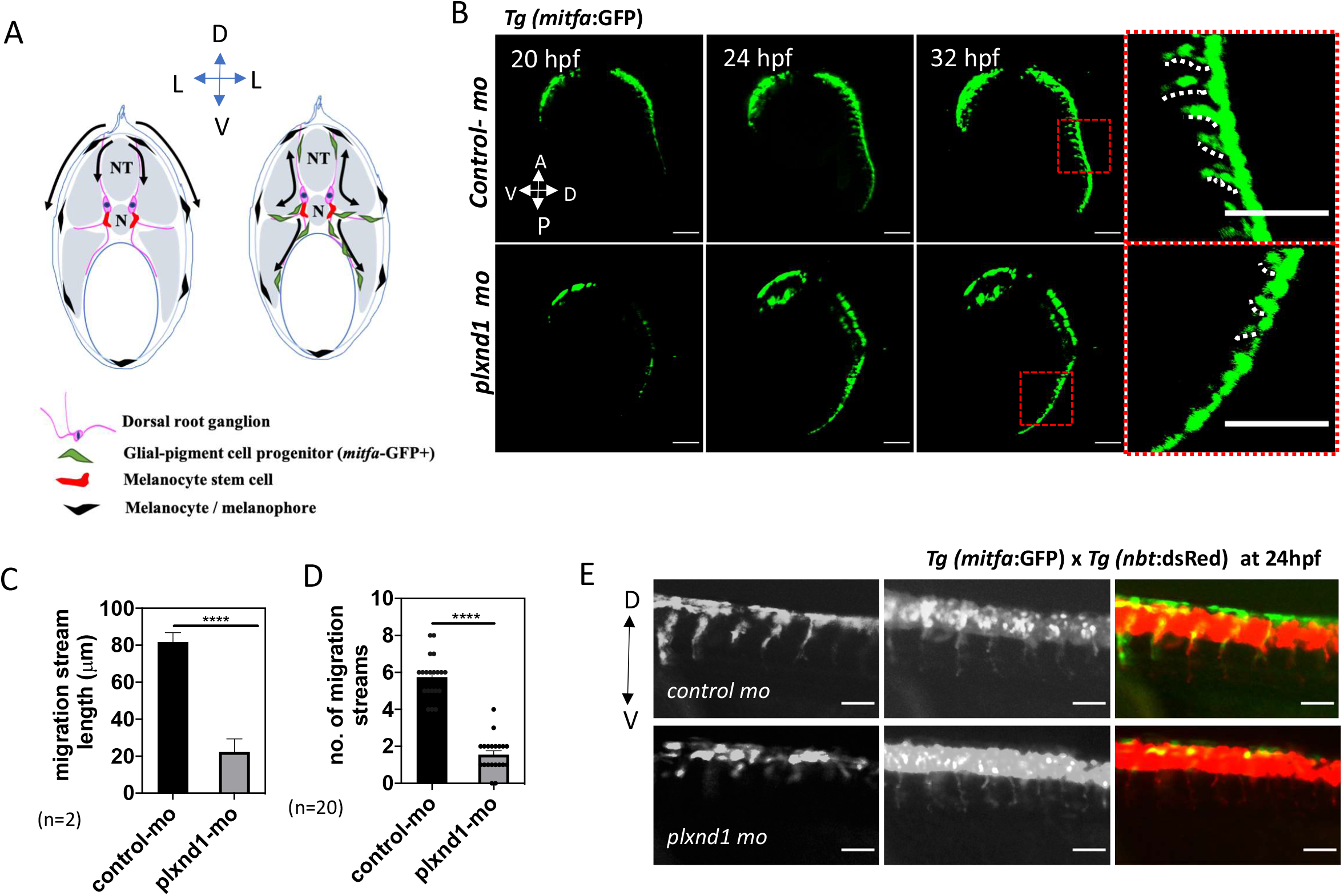
Loss of plxnd1 leads to delay in melanoblast streaming. (A) Schematic of (left) migration route along dorso-lateral and dorso-ventral axis, (on right) migration route along medio-lateral route, cell types are labelled at the bottom. (B) Snapshots at 20hpf, 24hpf and 32hpf from time-lapse imaging of *Tg mitfa:GFP* morphants. *in-sets* depict the streams arising from dorsal end marked along the dotted white line. Scale - 200µm (C) Quantitation of number of streams in morphants. (D) quantitation of stream lengths originating at dorsal end.10-12 streams were analysed from 3 animals per morpholino injection. error bar: mean ±s.e.m. **** represents a p-value <0.005. (E) Melanocyte streams marked with *mitfa:GFP* migrating along neuronal projections marked with *Tg nbt:dsRed*. Scale - 50µm

Collective streaming of NCCs requires co-polarization events. In the absence of guidance cues, cells would migrate individually but lack the invasiveness to enable colonization of NC derivatives in target tissue. Even at 20 hpf such migration defects were clearly manifested in *plxnd1* morphant *Tg (sox10: GFP*) animals (Fig S2A), raising a possibility that *plxnd1* may play a role in migration and establishment of other neural crest derived lineages as well. To investigate this we silenced *plxnd1* in other transgenic lines and studied the integrity of Sox10 derived enteric ganglia *Tg* (*nbt:DsRed*) and jaw cartilage *Tg* (*sox10:GFP*) at 5dpf. While we could not trace individual migratory events for these two lineages, we observed that these sox10-progenitor derived neural crest lineages were structurally intact in *plxnd1* morphants (Fig S2 B-D). These results suggested that role of plxnd1 in the positioning of NCC derived cells across the embryo may be selective to the melanocyte lineage.

Melanocyte precursors were earlier demonstrated to migrate along the projections of peripheral motor neurons [19]. Plexin-D1acts as axonal guidance receptor in several neurons and can thereby affect neuronal projections [26]. Therefore, we analyzed the patterning of melanocyte streams along the peripheral neuronal projection. For this, we used double transgenic zebrafish lines: *Tg (nbt:DsRed*) marking neuronal cells and *Tg (mitfa:GFP*) marking melanocytes, to discern the effects of plxnd1 loss on the ability of melanocyte precursors to migrate along peripheral neurons of the dorsal root ganglia. Interestingly, despite there being prominent axonal projections arising from peripheral neurons towards the ventral portion of the embryo, upon silencing of *plxnd1* melanocyte precursors fail to form migrating streams (Fig 2E) indicating a possible cell autonomous role of *plxnd1* in melanocyte migration. Further delving into the functional effects of PLXND1 signaling in melanocyte migration we analyzed the cytoskeletal organization and migration properties of cultured mouse B16 melanocytes upon activation or inhibition of PLXND1 mediated signaling.

### PLXND1 receptor mediates SEMA3E induced F-actin collapse and modulates directionality of migrating melanocytes

To investigate the role of Plexin-D1 it was important to identify its cognate ligand in the context of melanocyte functions. Plexin-D1can bind to multiple class 3 semaphorins as well as a class 4 member - SEMA4A. In all these cases it can couple to cytoskeletal changes by activating small GTPases *via* the GTPase activating protein (GAP) domain of the receptor [27]. Several studies have demonstrated its role either as an attractant or repellant in a context dependent manner (ligand and co-receptor) [26]. We identified the cognate ligand of *plxnd1* based on anti-sense morpholino based silencing of known *plxnd1* ligands in zebrafish. The ligand function was scored based on phenotypic absence of melanocytes along the lateral line in zebrafish embryos. Since semaphorins *sema3c* and *sema4a* had no apparent effect on embryonic pigment pattern development, the interpretation was inconclusive. However, silencing of *sema3e* recapitulated the lateral line melanocyte patterning in the embryos and suggested this to be the possible ligand involved in *plxnd1* activation (Supplementary Fig S3A & B).

As we had initially observed alterations in the expression of *Plxnd1* in mouse B16 melanoma derived cells, further experiments were carried out in these cells and later validated key observations in primary human melanocytes to delineate the cellular signaling effects. Treatment of B16 mouse melanoma-derived melanocytes with recombinant mouse SEMA3E resulted in immediate collapse in cell shape. This was clear upon staining the actin with phalloidin red to mark long filaments at the cell periphery as well as small actin foci (Fig 3A). SEMA3E treatment increased number of actin foci formation and concomitantly reduced the cell size (Fig 3B & C). Knockdown of Plexin-D1 (shPD) but not a non-targeting shRNA stable cells (shNT) inhibited SEMA3E induced collapse in cell shape by preventing the disassembly of filamentous actin (F-actin) (Supplementary Fig S3C-E). As plexins have structural/functional role in ECM attachment in cooperation with neuropilins, it was necessary to investigate whether F-actin disassembly is brought about by the PLXND1-SEMA3E signaling. Blockage of SEMA3E action by pre-treatment of cells with antibody against sema-binding domain of PLXND1 receptor (PD1 AB) but not normal rabbit IgG (IgG) prevented the cell collapse (Fig 3D top & bottom).

**Fig 3:**
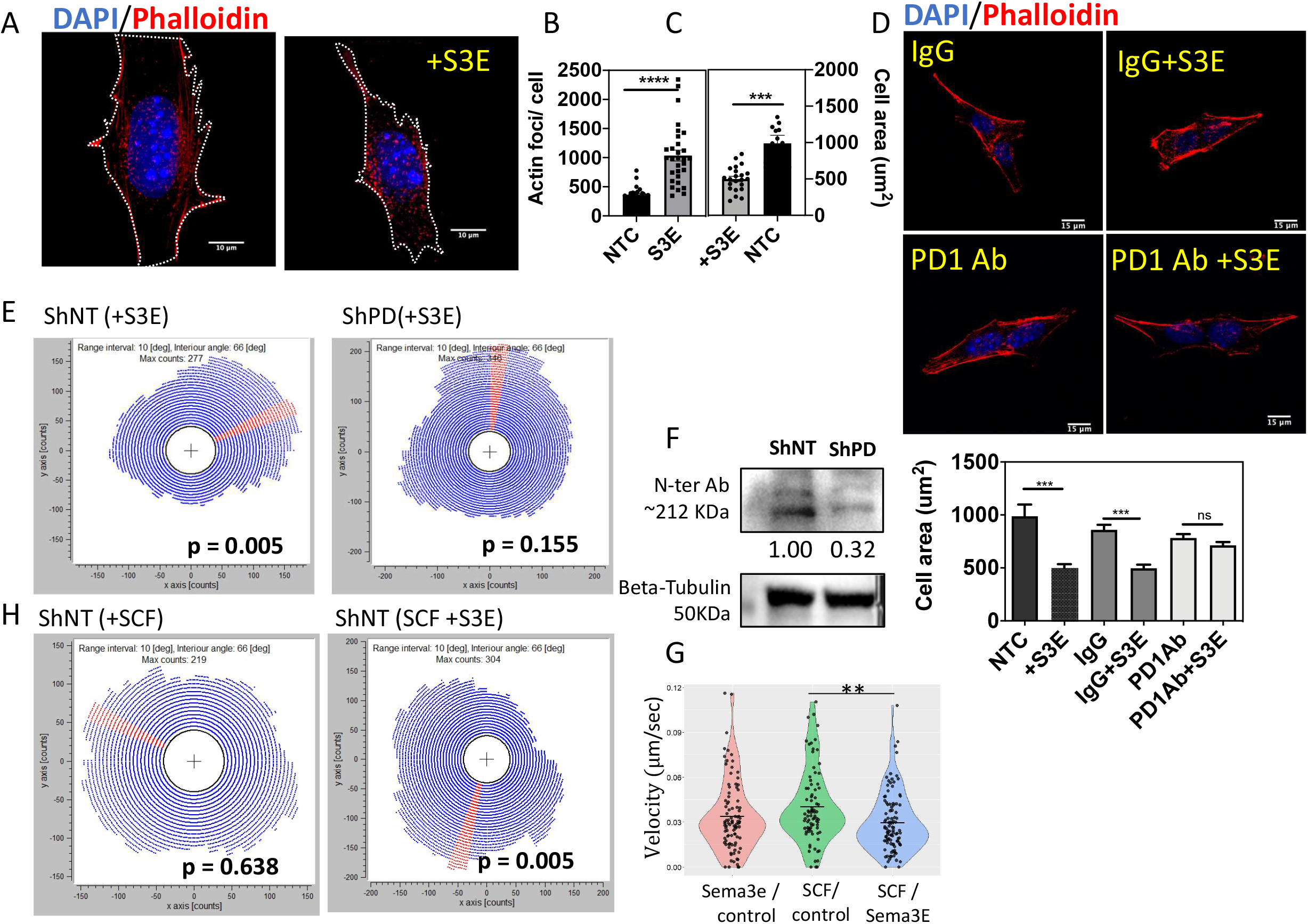
PLXND1 receptor mediates SEMA3E triggered F-actin collapse and affects directionality of migrating melanocytes. (A) Control B16 melanocytes or SEMA3E treated cells labelled with Phalloidin labelling F-actin, Scale bar=10 µm (B) Quantitation of nucleated actin foci (calculated as particles with aspect ratio = 1 and radius < 5µm) in control media or SEMA3E treated cells. (C) Quantitation of cell surface area of B16 melanocytes upon treatment with SEMA3E. For B & C: n>100 from 3 biological replicates represented as scatter plot. Two tailed t-test, p-value<0.001 is represented as***. (D) Phalloidin labelled B16 cell pre-treated with PLXND1 neutralizing antibody (PD1 Ab) or normal rabbit IgG as control followed by treatment with 5nM SEMA3E (+S3E), Scale bar=15µm, bottom: Quantitation of cell surface area upon antibody based neuralization of SEMA3E treated B16 cells. N=25 cells, 2 biological replicates. bars: mean±sem across 3 biological replicates. One way ANOVA Turkey’s multiple comparison test ***indicate P-value<0.05. (E) Rose plots depicting spatial distribution of shRNA either shNT non-targetting control or Plexind1 shRNA (shPD) transfected cells in chemotaxis chamber assay with treatments as indicated. In the circular rose plot, blue dots represent the angle position with the maxima of counts marked in red depicting the dominant direction of the cell motion across the gradient. P value calculated by Rayleigh’s statistic using Chemotaxis and migration analysis tool. (F) protein level of PLXND1 in shRNA expressing B16 cells shNT (non-targeting control) and shPD (plxnd1 targeting shRNA), numbers represent tubulin normalized fold change with respect to shNT. (G) Bean plot indicating the average track velocities across the experiment conditions. One way ANOVA, Turkey’s multiple comparison test, p value = 0.2742 non-significant. Bartlett’s stastistic P value=0.154. (H) Rose plots depicting spatial distribution of B16 cells in chemotaxis chamber assay in presence of SCF or in a gradient generated between SCF on one side and SEMA3E on the opposite side. In the circular rose plot, blue dots represent the angle position with the maxima of counts marked in red depicting the dominant direction of the cell motion across the gradient. P value calculated for Rayleigh’s statistic in Chemotaxis and migration analysis tool.

These morphological changes in cytoskeleton and cell shape are likely to have a direct consequence on cell motility. To understand the effect of SEMA3E-PLXND1 signaling on melanocyte migration we conducted a series of *in-vitro* chemotaxis assays. Time-lapse imaging (∼16h) of B16 cells subjected to a stable gradient of SEMA3E demonstrated the guidance mechanism. Control B16 cells expressing non-targeting shRNA (shNT) demonstrated that cells migrated away from the SEMA3E gradient. Thereby this signaling demonstrated a chemo-repulsive behavior in melanocytes. Whereas cells stably expressing shRNA against PLXND1 (shPD) failed to be repulsed by the addition of SEMA3E (+S3E) and migrated with reduced directional correlation (Fig 3E). Reduction in the protein level of PLXND1 could be captured by western blot analysis (Fig 3F).

Recent studies have demonstrated a combinatorial effect that various chemokine/chemotactic signals impose on migrating cells [28-30]. Studies on protective hyperpigmentation during the natural process of wound healing or epidermal tanning indicated that melanocytes proliferate and migrate towards cells secreting stem cell factor (SCF)[31]. Basal skin keratinocytes of mice engineered to express SCF recruit melanocytes in the epidermis and consequently have epidermal pigmentation [32, 33]. The ortholog of SCF in zebrafish kitlg is known to promote melanocyte migration and establish the McSC stem cell pool to result in lateral line melanophore formation [34, 35]. Thereby, we monitored cell migration by subjecting the cells to a combination of SCF and SEMA3E gradients.

Melanocytes demonstrated chemokinetic behavior towards SCF when provided in isolation and an enhanced velocity of migrating melanocytes could be observed (Fig 3G). In combination with SCF, SEMA3E caused a significant decrease in melanocyte motility, when compared against the SCF gradient alone (Fig 3H and Supplementary Table 2). However, the results indicated that SEMA3E driven guidance added a significant component of directionality to cells migrating in response to SCF. Primary human epidermal melanocytes behaved similarly and showed a chemo-repulsive behavior with respect to SEMA3E (Supplementary Fig S3F). Polarized migration is responsible for collective migration of melanocyte precursors as observed during the melanoblast streaming in dorso-ventral pathway [36, 37]. Combined with the observations on melanocyte migration in zebrafish embryos, SEMA3E-PLXND1 signaling pathway has the ability to guide melanocytes during embryonic pattern formation by channelizing melanocytes towards other known chemotactic cues such as SCF.

### SEMA3E-PLXND1 co-regulate melanocyte maturation program

Melanocyte fate establishment is fluid until completion of migration and differentiation. Early melanocyte lineage marker (*mitfa*) appears around 18h of fertilization in zebrafish embryo at dorsal end of neural tube in a rostro-caudal manner [38, 39]. By 24 hpf both lateral and medial streams are visibly migrating to populate the embryonic skin. Positional availability of chemotactic/chemokinetic cues in lateral and medial regions are likely to be independent and could lead to enforcement of distinct differentiation states. SEMA3E-PLXND1 signaling has an ability to activate intrinsic GAP activity and acutely remodel cellular cytoskeleton. While these effects are well-characterized, more recently the transactivating ability of PLXND1 on receptor tyrosine kinases has been delineated and complex interplay with other pathways is emerging [27, 40, 41]. The molecular responses of SEMA3E binding to PLXND1 receptor can lead to transcriptional regulation necessary for melanocyte survival and differentiation pathways.

Since the transcriptional responses could either be transient (like F-actin remodeling) or persistent, we treated B16 mouse melanocytes with recombinant mouse SEMA3E (5nM) for either 2h or 8h to capture both early and late responses. To distinctly identify transcriptional outcomes of signaling events from those elicited due to F-actin collapse, we compared them to signatures of Cytochalasin D (1µM for 8h) treatment. Early responses to SEMA3E treatment had several distinct transcriptional signatures associated with biological processes such as actin bundle assembly, cellular morphogenesis and GTPase signaling (Fig 4A). Interestingly *Gnas*, a gene that codes for Gs-α component of heterotrimeric G protein associated with cAMP signaling was found significantly upregulated upon 2h treatment with SEMA3E but not at 8h of treatment (Fig 4B). This gene is associated with pigmented *cafe-au-lait* skin spots and is a well-known pigmentation gene [42]. Pigment synthesis in differentiated melanocytes is dependent on cAMP mediated signaling and is controlled by melanocyte stimulating hormone œ-MSH by activating the MC1R receptor which couples with the adenylate cyclase using GNAS [43, 44]. We therefore hypothesized that SEMA3E treatment could couple activation by œ-MSH.

**Fig 4:**
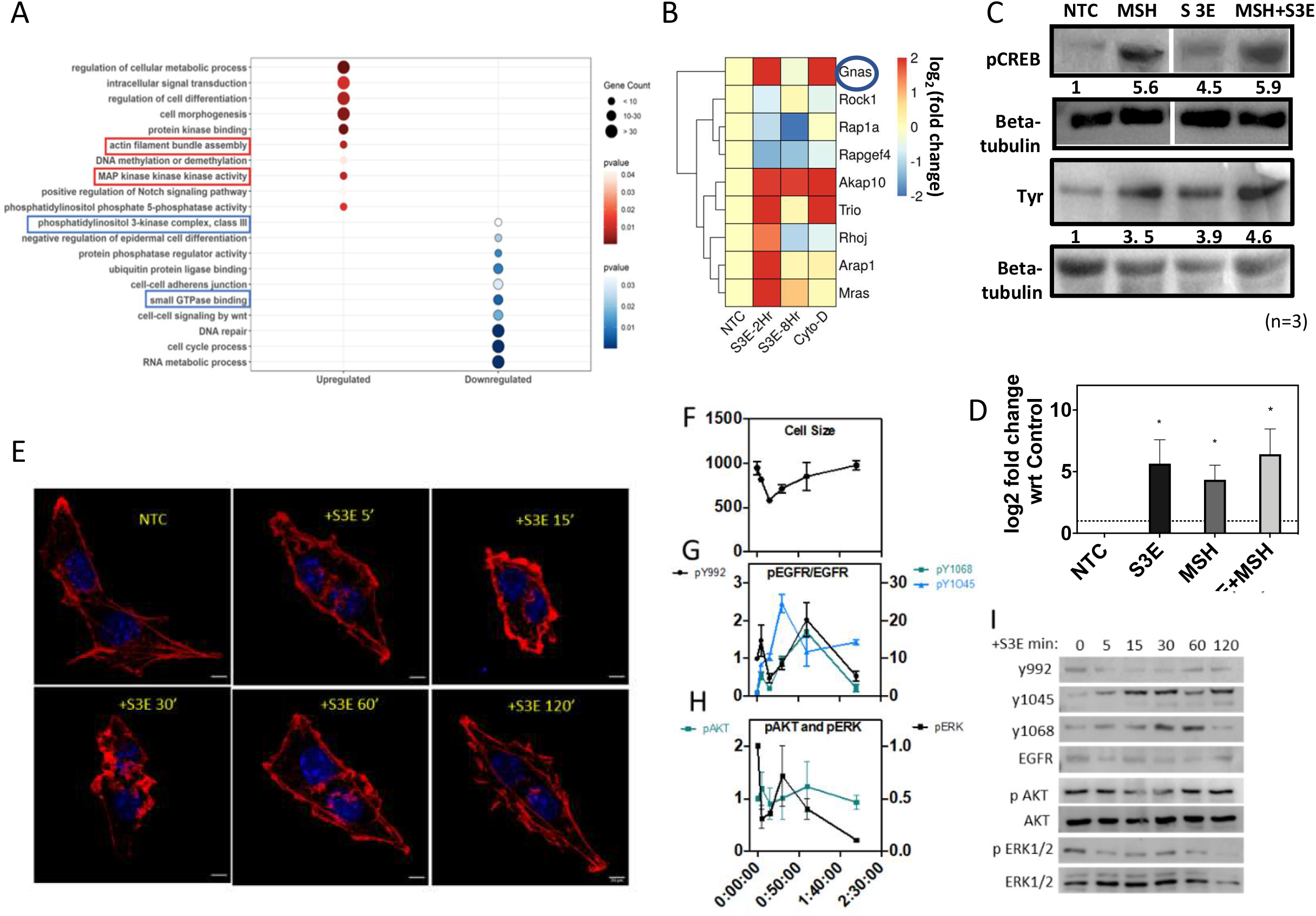
Gene expression changes and melanocyte functioning upon Sema3e treatment. (A) Biological processes enriched upon SEMA3E treatment with an adjusted p value < 0.05 (upregulated in red, downregulated in blue) size of the circle depicts the number of differentially regulated genes with a p value of < 0.05. (B) mRNA expression of GTPases binding proteins (with a p value of < 0.05) depicted as heatmap, that are enriched upon SEMA3E treatment. (C) Levels of pigmentation associated proteins expressed in B16 cells upon treatment with SEMA3E (n=2 replicates). Values represent β tubulin normalized fold change over no treatment control (NTC). (D) Quantitation of *Mitf* expression mRNA levels upon SEMA3E treatment across two biological replicates. (E) Changes in cell shape upon treatment with SEMA3E analyzed by estimating area marked by phalloidin staining of F-actin, scale=20µm (F) Comparative trends of a cell shape changes and (G) protein phosphorylation states of EGFR, (H) AKT and ERK1/2. Values represent mean±SEM across biological triplicates. (I) Representative western blot analysis of dynamic regulation of protein kinase phosphorylation states upon various durations of SEMA3E treatment.

Treatment of SEMA3E is sufficient to elevate CREB phosphorylation and induction of tyrosinase perhaps due to the elevated basal activity of MC1R. Pretreatment of B16 cells with SEMA3E followed by MSH addition resulted in a further increase in CREB-phosphorylation (Ser133) and tyrosinase levels indicating synergism (Fig 4C). An induction in the *Mitf* mRNA levels could also be observed with SEMA3E treatment (Fig 4D). These results suggested a role of SEMA3E-PLXND1 signaling in amplifying the process of melanocyte maturation by pre-conditioning the cells for α-MSH mediated activation of the differentiation program.

To map the transcriptional responses of SEMA3E to other pathways we performed connectivity map (CMAP) analysis using CLUE platform developed by Broad Institute, USA and compared the gene expression profile elicited by treatment of SEMA3E in melanocytes to other existing melanocyte datasets (Fig S4). One of the key pathways that emerged was the EGFR signaling pathway. Earlier study by [40] had demonstrated a physical association between PLXND1 receptor and ERBB2 and implicated a role in melanoma metastasis. Role of ERBB3B receptor that signals via the EGFR pathway in the establishment of melanocyte stem cell pool is well known [45]. Therefore, we analyzed the temporal effect of SEMA3E mediated stimulation on different EGFR phosphorylations (Fig 4 E-I): Y992 involved in PLCγ-mediated downstream signaling, Y1045 involved in receptor ubiquitination and degradation) and Y1068 associated with GRB2 mediated signaling. SEMA3E treatment showed a marginal increase in Y992 phosphorylation, with a peak around 60 min of treatment. Whereas changes in the phosphorylation at Y1045 and Y1068 were robust. Y1045 phosphorylation was activated within 5 min of SEMA3E stimulation and continued throughout the treatment duration. EGFR phosphorylation at Y1068 progressively increased and peaked between 30 min to 60 min of SEMA3E treatment before reverting to basal level. Changes in cell surface area as indicated by F-actin staining corresponded with changes in EGFR phosphorylation at Y992 – an initial collapse followed by gradual recovery of cell surface area (Fig 4E &F). The status of ERK1/2 phosphorylation remained below the basal levels (Fig 4H). AKT phosphorylation on the other hand remained constant. These observations suggested SEMA3E-PLXND1 signaling has an acute and significant effect on EGFR signaling events and could regulate melanocyte fate transitions during early development that are downstream of this signaling pathway.

### sema3e-plxnd1 signaling establishes melanocyte stem cells for pigment patterning

Zebrafish mutants for *erbb3b* gene (*picasso*) develop normal pigment pattern during embryonic stages but have a significant loss in melanocytes that arise from stem cell pool during and after metamorphosis. We have so far demonstrated altered motility of melanocyte precursors and defective positioning of melanocyte stem cells. To explore the status of melanocyte stem cells that go on to populate adult animals, we treated *plxnd1* morphants at 24hpf with a melano-toxic chemical 4-(4-Morpholinobutylthio) phenol (MoTP) until 72hpf (Fig 5A). Once melanocytes were ablated these animals were transferred to fresh medium to allow melanocyte regeneration. At 72hpf when the morphants were transferred to fresh water post-MoTP treatment, the pigmented melanophores were completely ablated in both plxnd1 morphants and control morphants [45]. Upon careful examination, positioning of cells with GFP expression in *Tg mitfa:GFP* cells could be clearly visible as MoTP resistant GFP positive cells in control animals, *plxnd1* morphants demonstrated absence of GFP positive MSCs in the lateral line (Fig 5B). This could be due to a decreased mobility of melanocyte precursors during 24-48 hpf resulting in positioning of these precursors in alternate areas in the embryo. Since we did not observe extraneous melanophores, it is likely that these cells are programmed to cell death. Substantiating this possibility, the frequency of tunnel-TMR stained cells in *plxnd1* morphants at 36 hpf was found to be higher near the head and trunk region, indicating a greater amount of cell death (Fig 5C).

**Fig 5:**
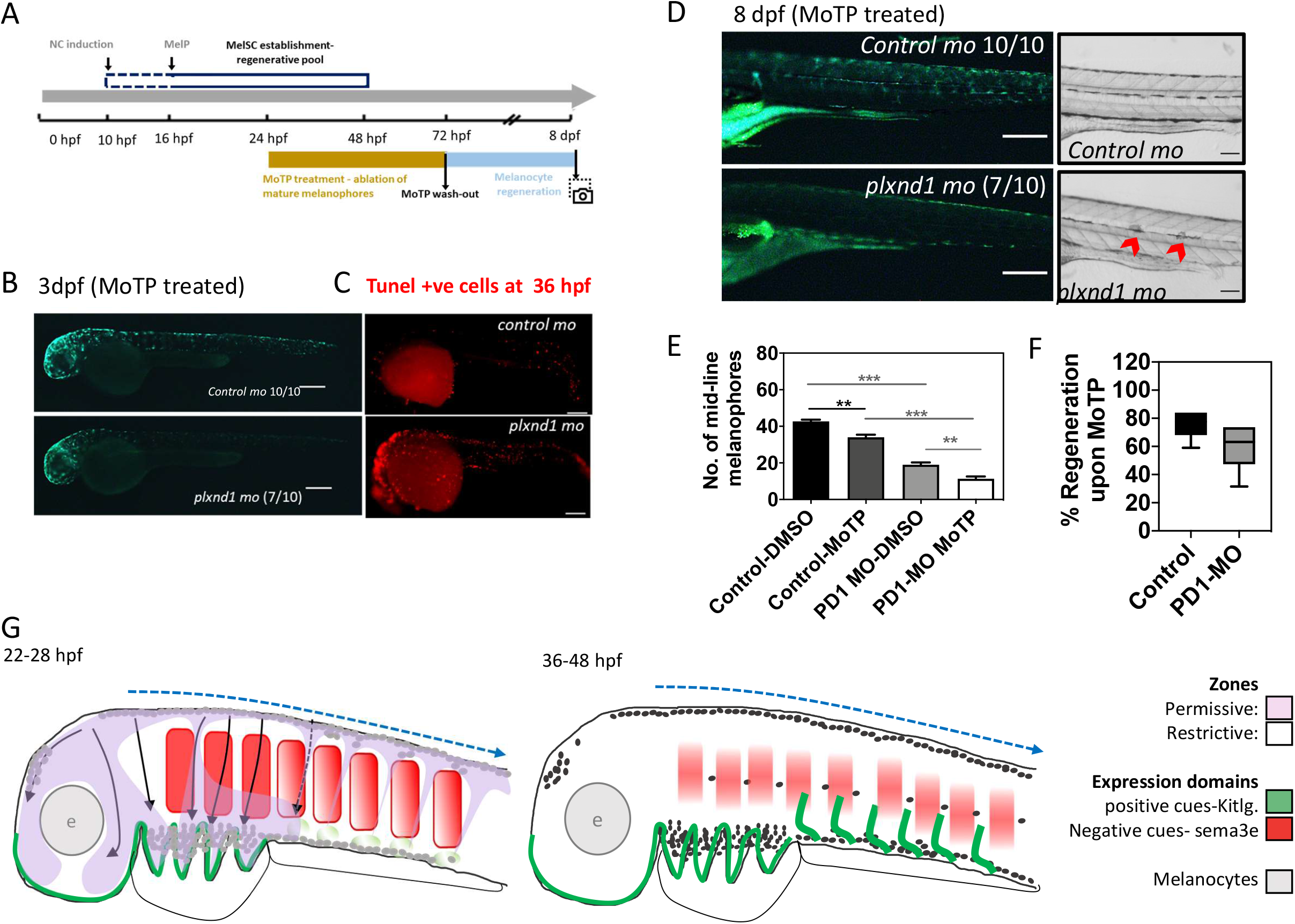
plxnd1 is essential for maintenance of melanocyte stem cell pool and regeneration. (A) Experimental strategy for MoTP (4-(4-Morpholinobutylthio) phenol) based melanocyte ablation and subsequent regeneration in morphants. (B) Stitched images of the zebrafish embryos expressing *GFP* under *mitfa* promoter upon completion of MoTP based melanophore ablation at 3dpf. Scale 30µm. (C) Representative images of tunnel-TMR stained lateral trunk region in morphants indicating extent of apoptosis at 36 hpf. Scale 30µm. (D) lateral trunk region with regenerated melanophores at 8 dpf in MoTP treated animals. Scale 100µm. (E) Quantitation of mid-line melanophores in morphants at 8dpf (post-regeneration) in MoTP treated morphants. Turkey’s Multiple Comparison Test, ** p value <0.05, *** p-value <0.005. (F) Melanophore regeneration represented as a percent of regeneration *wrt* corresponding control or plxnd1 morphant, unablated embryos in MoTP treated. Unpaired T-test, p value<0.06. (G) Graphical presentation of the model for melanophore migration and mid-line melanophore establishment. Relative chemotactic gradients and melanophore positioning during melanocyte pattern development in zebrafish.

Upon completion of regeneration at 8dpf, control morphants animals regenerated melanocytes to around 80% or more, however *plxnd1* morphants had a lower regeneration potential of less than 60% (Fig 5E&F). While, it was difficult to estimate cell death specific to melanoblast, a clear decrease in *mitfa:GFP* tagged cells in MoTP treated *plxnd1* morphants at 72hpf and lower pigmented regenerated melanocytes at 8dpf indicated towards a possible event of melanoblast death in *plxnd1* morphants.

## Discussion

Self-assembly of characteristic patterns has been a matter of utmost fascination to biologists. To define and reconstruct pigment patterns, a comprehensive understanding of the involved players is necessary. Positive cues such as SCF that enable melanocyte migration are identified and well-characterized across several model systems, whereas the known negative cues so far happen to be contact dependent. Hence identification of a signaling pathway that restricts the migratory path and enables directionality, is an important milestone in our understanding of the behavior of neural crest derived cells. So far, such exquisite guidance mechanisms involving localized grandient of inhibitory cues have not been reported for melanocytes.

The path taken by melanocytes is distinct and well known. Knowledge of various positive cues and negative contact dependent signals explain the migratory path to a large extent. However, migration as distinct organized streams from specific dorsal locations remained perplexing. Based on our understanding of the role of plexin D1and semaphorin 3E deciphered here, and earlier studies on the expression pattern of semaphorin 3E and kitlg (SCF) [34, 46, 47], we propose that melanocyte migration path is governed by the zone of expression of these two guidance factors (Fig 5G).SEMA3E-PLXND1 signaling pathway is critical for restricting melanocytes into orchestrated streams for targeting melanocytes to dermal regions towards SCF expression zones. It is likely that the integration of cues on melanocyte decides the choice of migratory path and is observable as directional consolidation in chemotactic migrations observed *in vitro* against SCF and sema3e gradient contrasts.

Pathways that promote migration, such as SCF and endothelins have almost always been found to have a pro-melanocyte fate biasing effect as they impinge on the MITF gene-regulatory network. Hence, our understanding of these pathways in migration and fate determination have remained intertwined. Contact dependent restrictive cues primarily control migration and positioning of melanocyte, and their effect if any on melanocyte gene expression remain unexplored. Interestingly, the SEMA3E-PLXND1 guidance cue also impinges on the core pathways of melanocyte stem cell establishment and differentiation by co-opting other signaling pathways. While the transactivation of EGFR is likely due to direct interaction mediated activation, potentiation of the melanocortin pathway is indirect, *via* the expression of the MC1R effector GNAS. The latter effect is an outcome of cell shape change brought about by disruption of the F-actin filaments, as this is recapitulated upon cytochalasin D treatment.

Guidance mechanisms have a transient yet profound implications on cellular fate and functioning. This could be evident in the establishment of lateral line melanophore forming, regeneration competent stem cells. Previous reports have reported expression of PLXND1 in invasive melanoma tissues, suggesting a role in cancer metastasis [40]. Given the recapitulation of neural crest like characteristics in melanoma, it is likely that the invasive properties may arise from an expression pattern reminiscent of earlier developmental stages. This understanding of melanocyte responses to SEMA3E-PLXND1 guidance signaling would in future pave the path to predictably influence migration in cancerous conditions such as melanoma and in repopulating melanocytes in degenerative disorders such as vitiligo.

## Methods

### Zebrafish maintenance

The zebrafish lines used in this study: Assam wildtype - ASWT, *Tg(−4*.*9 sox10:egfp)ba2, Tg(mitfa:GFP), Tg(ftyrp1:GFP), Tg(Xla*.*Tubb:DsRED)* and double labelled *Tg(Fli:GFP/Gata:DsRed)* (details and references are provided in **Supplementary Table 1**). The zebrafish lines were bred, raised, and maintained as described in (Westerfield, 2000). Embryos were staged according to (Kimmel *et al*., 1995). Embryos older than 24 hpf were treated with 0.003% PTU (1-phenyl-2-thiourea) to inhibit pigment formation whenever required, to aid in fluorescence imaging. Zebrafish handling and experiments were performed in accordance with the protocols that were approved by the Institutional Animal Ethics Committee (IAEC) of the CSIR-Institute of Genomics and Integrative Biology, India under the rules and regulations set by the Committee for the Purpose of Control and Supervision of Experiments on Animals (CPCSEA), Ministry of Environment, Forests and Climate Change, Government of India.

### Fluorescence activated cell sorting, RNA sample preparation and Microarray

The microarray-based gene expression analysis was conducted in association with a senior colleague and the experimental details are in greater detail described in previously submitted thesis. Briefly, the fluorescently labelled cells were isolated by FACS from single cell suspensions made out of *Tg(mitfa:GFP)* embryos during different stages of melanocytes development - 18-22 hpf melanoblasts (bMEL in dorso-lateral streams), 28-32hpf immature melanocytes (iMEL) and 3dpf mature pigmented melanocytes (mMEL, mature). FACS was performed using BD Aria III. RNA was isolated from each of these sorted populations according to manufacturer’s protocols (Nucleospin RNA XS kit; MachereyNagel). After ascertaining the quality by BioAnalyser and scoring for enrichment of melanocyte specific genes by quantitative real-time PCR the samples were sent for gene expression analysis by microarrays at Genotypic technology, Bangalore, India. The analysis was performed using Genespring GX software. Microarray analysis was performed using Zebrafish_GXP_8X60K (AMADID: 74191) array slides using T7 promoter-based linear amplification to generate labelled complementary RNA. Manual image quality control for the slides was performed and found to be devoid of uneven hybridisation, streaks, blobs and other artefacts. Intra array quality check and percentile shift normalisation were performed using GeneSpring GX software. A (−0.6 to 0.6) log_2_ fold change cut off was employed to ascertain differentially expressed genes in this dataset and p-value ≤ 0.05 to determine the consistency across biological replicates.

### Percoll based density centrifugation of pigmented melanophores

A modified protocol from Yamanaka H *et al* [48] was used to isolate pigmented melanophores from 3dpf zebrafish embryo. Briefly, the embryos were anesthesized and their heads were cut-off using scalpel to separate out pigmented retinal epithelial cells (RPE). Embryo bodies were then incubated in TrypLE solution (Gibco 12604013) with occasional pipetting to facilitate formation of cell suspension. The cell suspension was filtrated using a 22-μm mesh and then gradient-centrifuged with 50% Percoll (Sigma) at 30 × *g* and 28 °C. The pellet was resuspended in serum-free L15 (Gibco) medium. Cells were then washed with DPBS and immediately processed for RNA isolation according to manufacturer’s protocols (Nucleospin RNA XS kit; MachereyNagel).

### Anti-sense Morpholino based gene silencing

All morpholinos (MOs) designed in this study were obtained from GeneTools. For preliminary screen for plexin receptors involved in melanocyte guidance one translational block mopholino each were against *plxn b1a, plxn b1b, plxn c1* and *plxn d1*. To validate the role of plexin d1 that emerged as significant receptor in melanocyte patterning from preliminary screen a previously reported splice-block morpholino against *plxnd1* (*plxnd1* SB-MO) was used. 3 other morpholinos individually targeting the cognate plxnd1ligands – sema3g, sema3e and sema4d were designed for identifying the ligand involved in Plxnd1 mediated guidance response in zebrafish melanocytes during development. Standard control MO and p53 MO was obtained from GeneTools. Morpholinos were designed using gene tools website (http://www.gene-tools.com/).

### Imaging

Bright field imaging of live embryos was performed using Zeiss (Stemi 2000C) or Nikon smz800n. For initial phenotyping based on counting of pigmented melanocytes in brightfield, the embryos were treated with 0.003 % PTU beginning at 20 hpf until 3 dpf and then transferred to plain E3 water to initialize pigment synthesis. This allowed resolution between 2 adjacent melanophores that are otherwise difficult to count as separate cells. Zeiss axioscope A1 microscope (with AxiocamHRc) was used for imaging of fluorescent embryos. Images were further processed and analyzed in Adobe Photoshop CS3 or Fiji. The embryos were either treated with tricaine or embedded in 1.5 – 2% of methylcellulose to restrict their movement during imaging.

### Time-lapse imaging of zebrafish embryos

Live imaging of *Tg(mitfa:GFP)* embryos between 28 hpf-36 hpf were done using Leica SP8 STED confocal microscope. Images were captured in xyzt mode at 10x magnification. 40 mm tissue culture dishes (Orange scientific) were used, on this the embryo was placed in a drop of E3 water and then overlaid with 2% Low melting agarose (LMA). The embryo was laid laterally to visualize dorso-lateral cell migration in trunk region. The plate was then filled with E3 water, and the humidifier was used when the temperature was set at 320C so as the temperature of the E3 water was maintained at 26-28 °C. The images were sequentially captured first in green channel (488nm) and then in transmitted light. An interval of 30 minutes was maintained between each acquisition. Experiment was then exported as TIFF images and video files were created in the quicktime mode. The images at each minute interval were taken up for analysis using ImageJ (Fiji).

### Cell culture

B16 mouse melanocytes (kind gift from the lab of Dr. Satyajit Rath at NII, New Delhi, India) were cultured in bicarbonate buffered DMEM-High Glucose (Sigma. D-5648), supplemented with 1x anti-anti (Gibco), 10% FBS (Gibco) at 37°c with 5% CO_2_.

NHEM neo-Normal human epidermal melanocytes (Lonza CC-250) were grown in proliferative conditions in PMA containing medium MBM4 (Lonza CC-3249). For differentiation, cells were switched to M254 medium for 3-4 population doublings (Thermofisher Scientific, Life Technologies).

### F-actin integrity assay

Cells cultured to a confluence of 50,000 cells per cm^2^ on coverslips were treated with 5nM purified recombinant mouse SEMA3E (R&D biosystems, catalog no. 3238-S3-025) for 15 mins at 37°C. Immediately after the incubation the cells were washed with 1x DPBS, twice and then fixed with 4% paraformaldehyde at 37°C. Once fixed the cell were either stored in DPBS at 4°C or taken ahead for F-actin staining with Phalloidin (Alexa Fluor 568 Phalloidin-A22283, ThermoFisher scientific). Phalloidin stained coverslips were then imaged at high magnification with GE DeltaVision 100x oil immersion objective. Cell surface area was measured using Fiji analysis by marking the cell shape in brightfield and actin foci were calculated by the measure particles function in fiji after setting the colour threshold for the red channel. The particles were estimated based on set particle diameter of 0.1 μm diameter and aspect ratio of 1.

### Chemotaxis assays

Chemotaxis chamber assay was set-up in according to the protocol described by the manufacturer. Briefly, each chemotaxis coverslip has 3 (1mm wide) trough regions where the cells are seeded. Each trough is surrounded by media reservoirs on either side. The trough is connected to media reservoirs on either side by a small opening that allows for gradual diffusion of chemotactic agent across the trough, generating a concentration gradient. Once the cells are adhered and the chemotactic agent(s) are added into the respective reservoirs the chamber is set-up on the microscope stage for imaging. Growth promoting chemokine stem cell factor (SCF, PeproTech #300-07) was used for competition against SEMA3E. Time-lapse XYT imaging was done for ∼20 hours with time interval of 30mins. During the imaging the slide was maintained at 37°C with 5% CO_2_ using Environment controlling unit. Imaging was done with GE-DeltaVision at 10x magnification. Chemotaxis analysis was done by individually generating the tracks of each cell (>100 cell/per experiment) using manual tracking plugin in Fiji and then processing the track information in chemotaxis and migration tool (Fiji Plugin) developed by ibidi.

### SEMA3E neutralization assay

The cells were pre-incubated with different concentrations of anti-N-ter-PLXND1 antibody (anti-NPD1 antibody) for 2 hours at 37°C. After this the cell were treated with recombinant SEMA3E and analysed for F-actin stability

### Microarray sample preparation from treated B16 cells

B16 cells were cultured in 25cm^2^ tissue culture flasks and treated with 5nM recombinant SEMA3E (R&D biosystems, catalog no. 3238-S3-025) for either 2 hours or 8 hours. To distinctly identify transcriptional signatures unique to loss of actin integrity, the cells were treated with 1μg/ml Cytochalasin D (1mglml stock solution in DMSO. C8273, Sigma-Aldrich) for 8 hours. RNA was isolated according to manufacturer’s protocols (Nucleospin Triprep kit; MachereyNagel). Isolated RNAs were analyzed using BioAnalyzer and after satisfactory results microarray was performed on Agilent Mouse gene expression microarrays at Genotypic technology, Bangalore, India. All quality control measures were taken before executing the microarray. Mouse GXP_8X60k AMADID: 065570 slides were used for microarray and Agilent Quick-Amp labelling kit (p/n5190-0442; T7 promoter-based linear amplification to generate labelled complementary RNA). All QC and further pipeline are similar as described in section 2 of methods. Data pertaining both the array experiments have been deposited in Gene Expression Omnibus.

### Melanocyte regeneration model establishment

*plxnd1* morphants and control morphants were dechorionated at 24 hpf and treated with 100 μM MoTP till 72 hpf to ablate melanized melanophores. At 72 hpf, the embryos were rinsed twice with embryo water to remove residual MoTP. Treated embryos were grown in embryo water till 8 dpf and subsequently imaged to assess melanophore regeneration.

### In-situ Tunnel-TMR staining

Morphants between 32 hpf-48 hpf were dechorionated, anesthetized and fixed in ice-cold 4%PFA for 2h at room temperature. Tissue was then permeabilized in a graded ethanol series (50%, 70%, 95% and 100%) followed by 10 min incubation in acetone at -20°C. permeabilized embryos were washed thrice with PBS and then treated with Proteinase K (10 µg/µl) for 30 mins. After washing thrice with PBS the embryos were incubated in Tunnel-TMR labelling mix (In Situ Cell Death Detection Kit, TMR red, Cat. No. 12 156 792 910) for 1h at 37°C in a humidified condition. Once labelled the embryos were washed and stored in PBS until imaged.

### Connectivity Map analysis

A set of significantly up and down differentially regulated genes was used to query the connectivity map database (P-Value<0.05). Clue.io database was used to fetch the relevant drug treated expression profiles that were mimicking/reversing the SEMA3E treated gene expression signatures (https://clue.io/; dated ∼10.04.2020). A cut-off of +/- 90 connectivity score and its treatment in A375 cell line was applied to filter the drugs. The complete gene expression profile for all of the filtered drugs was fetched and genes were classified as up/down regulated upon a respective drug treatment if it had a Z-score of +/- 0.5.

### CRISPR mutant generation

Short guide RNA (sgRNA) targeting zebrafish *plxnd1* gene were selected from ECRISP 13 (http://www.e-crisp.org/E-CRISP/) online tool using default parameters. Primers were designed for generating the complete sgRNAs (supplementary table 1) using annealing PCR. The products were gel eluted and purified using Qiaquick Gel extraction kit (Qiagen Cat no. 1628704) according to manufacturer’s protocols. In vitro transcription was performed on these PCR products using Megashortscript kit (ThermoScientific®;AM1354) according to manufacturer’s protocol. These sgRNAs and Cas9 protein were (a kind gift from Dr. Souvik Maiti, CSIR-IGIB, India) incubated on 25°C to form complex and 100pg of sgRNA and 300pg of spCas9 protein was injected into single cell zebrafish embryos. The injected embryos were tested for IN/DEL using T7 endonuclease assay and confirmed by sequencing.

We adapted cell-type specific knockout stratergy using MinicoopR vector in which Mitfa promotor (melanocyte specific promotor) drives Cas9 and *mitfa* mini gene and has space for two sgRNA under Ubiquitous (U6) promotor. We replaced *mitfa* mini gene with GFP using restriction digestion approach. We injected mitfa :Cas9, mitfa :gfp plasmid a little ahead of one cell stage in zebrafish embryo and imaged it at 2dpf stage. Figure1H shows labelling of majority of melanophores.

We designed two sgRNA using CHOPCHOP (https://chopchop.cbu.uib.no/) targeting plxnd1 gene. The cloning was performed by digesting the plasmid using BseRI enzyme. We confirmed the clones using PCR based approach using Forward primer specific to plasmid and sgRNA as a reverse primer. The validated clones were injected at one-cell stage in Zebrafish embryos and the imaging was done at 3dpf stage.

## Competing Interests

Authors do not have any conflict of interest.

## Acknowledgements

This work was supported by the Department of Science and Technology supported the work through the grant (GAP165). Council of Scientific and Industrial Research (CSIR), India, through grant RegenX-MLP2008 and Department of Biotechnology through the grant (GAP0182) provided support to the execution. YSJ acknowledges DBT for Research Fellowship. We thank Dr Rajesh S Gokhale for the meaningful discussions and constant support throughout the execution of this work.

## Data Submission

Microarray Data pertaining to this work has been submitted to Gene Expression Omnibus repository **GSE189059** and **GSE189438**

## Supplementary Material

Supplementary Table 1

Details of the reagents and their source information is provided

Supplementary Table 2

Cell migration track information for B16 cells exposed sema3E, SCF and in combination.

Supplementary Video 1 and 2

Video files of melanocyte migration in developing control and plexin D1 morphant embryos

## Supplementary Figure Legends

**Fig S1: Identification of plxnd1 as cell surface receptor essential in melanocyte patterning**.

(A) Fold enrichment of Mitf expressing cells upon FACS sorting or density centrifugation. (B) Heatmap of log2 (fold change) in genes involved in melanocyte differentiation during melanocyte development. Size of the circle is proportional to the number of genes representing that pathway and the corresponding adjusted p value scales are depicted. (C) (right) Heatmap of log2 (fold change) in transcription factors involved in melanocyte specification during melanocyte development. (left) Heat-map of key cell adhesion and melanocyte differentiation genes upregulated during the McSc to pMel transition. (D) Lateral embryonic melanophore patterning in *plxnb1a, plxnb1b* and *plxnd1* (splice block) morphants at 5dpf, n=20. Scale-100µm. (E) Quantitation of mid-line melanophores in Morpholino against different plxns injected animals at 2dpf, n=25 each in 3 biological replicates. *** indicates a p-value < 0.001 and bottom bar graph depicts quantified lateral mid-line melanophores. (F) Western blot analysis of control and *plxnd1* morphant embryo (plxnd1 MO1) lysates probed with PLXND1 antibody and normalized to β-Actin. (G) Vascular branching in control or *plxnd1* morphants at 2dpf marked by fli:GFP, in-sets depicting branching defects at dorsal end. Scale bar=50µm. (H) F0 zebrafish embryos at 5 dpf injected either with Cas9 protein alone (top) or with Cas9 protein along with plxnd1 sgRNA targeting the *plxnd1* locus (bottom). Red arrowheads indicate lateral line melanophores. (I) Amplicon sequencing of F0 animals at the CRISPR locus in plexin D1.

**Fig S2: Decrease in PLXND1 levels leads to delay in melanoblast streaming**

(A) Migrating Sox10:GFP positive cells between 18-22hpf in morphants. (B) Positioning and integrity of Sox10 derived enteric ganglia. (C) Quantitation of number of enteric speckles in morphants. Two tailed t-test, p value=0.49, indicating the difference to be non-significant. Error bars= mean±SEM across two biological replicates, n=20. (D) Integrity of sox 10 derived Jaw cartilage in morphants. me: mesenteric cartilage, cb1-5: cerato-brachial arches, Ch-Cerato Hyoid. Scale bar = 100µm.

**Fig S3: PLXND1 receptor mediates SEMA3E triggered F-actin collapse and affects directionality of migrating melanocytes**

(A) Embryonic melanocyte patterning at 5dpf in morphants of cognate semaphorins that are known to function with plxnd1, in-sets depict the lateral trunk region with pigmented melanocytes, scale bars=200µm. Red arrowheads point out mid-line lateral melanophores. (B) Quantitation of melanocytes at lateral mid-line in semaphorin morphants. Error bar= mean±sem; n=20 per morpholino, 2 biological replicates. *** depicts p-value<0.01, one-way ANOVA, Dunnett’s multiple comparison test. (C) F-actin labelled with Phalloidin in shNT (non-targeting short hairpin) and shPD (short hairpin targeting plxnd1) expressing B16 cells treated with control media or SEMA3E, Scale bar=20 µm Average velocity of individual cell tracks during chemotaxis assay. Two tailed t-test, * represents p-value <0.0018. bars-mean±s.e.m. (E) Quantitation of nucleated actin foci in control media or SEMA3E treated shRNA expressing cells. (F) Rose plots depicting the spread of migrating primary human melanocytes in chemotaxis chamber assay upon exposure to chemotactic gradients. P value calculated by Rayleigh’s statistic in Chemotaxis and migration analysis. N>250 cell, 2 replicates. (G) Quantitation of cell surface area foci of control media or SEMA3E treated shRNA expressing B16 cells.

**Fig S4: Connectivity map analysis of SEMA3E treated gene expression data**

(A) Heatmap depicting the expression status of genes upon indicated drug treatment. The green-red colour of the heatmap are the Z-score values from connectivity map analysis. Here, Rows represents the drugs and column represents the genes (downregulated genes upon *SEMA3E* treatment). The top multi-coloured annotation bar represents the class of a gene and the right-hand side annotation bar plot represents the percentage of the gene that is getting up/down regulated upon the respective drug treatment (yellow represents down and blue represents up).

## Notes

### Competing Interest Statement

The authors have declared no competing interest.

